# Functional Improbable Antibody Mutations Critical for HIV Broadly Neutralizing Antibody Development

**DOI:** 10.1101/262592

**Authors:** Kevin Wiehe, Todd Bradley, R. Ryan Meyerhoff, Connor Hart, Wilton Williams, David Easterhoff, William J. Faison, Thomas B. Kepler, Kevin Saunders, S. Munir Alam, Mattia Bonsignori, Barton F. Haynes

**Affiliations:** Duke Human Vaccine Institute, Duke University, Durham NC 27710; Department of Microbiology and Immunology, Boston University, Boston, MA

## Abstract

HIV-1 broadly neutralizing antibodies (bnAbs) require high levels of activation-induced cytidine deaminase (AID) catalyzed somatic mutations for optimal neutralization potency. Probable mutations occur at sites of frequent AID activity, while improbable mutations occur where AID activity is infrequent. One bottleneck for induction of bnAbs is the evolution of viral envelopes (Envs) that can select bnAb B cell receptors (BCR) with improbable mutations. Here we define the probability of bnAb mutations and demonstrate the functional significance of key improbable mutations in three bnAb B cell lineages. We show that bnAbs are enriched for improbable mutations, implying their elicitation will be critical for successful vaccine induction of potent bnAb B cell lineages. We outline a mutation-guided vaccine strategy for identification of Envs that can select B cells with BCRs with key improbable mutations required for bnAb development. Our analysis suggests that through generations of viral escape, Env trimers evolved to hide in low probability regions of antibody sequence space.

## INTRODUCTION

The goal of HIV-1 vaccine development is the reproducible elicitation of potent, broadly neutralizing antibodies (bnAbs) that can protect against infection of transmitted/founder (TF) viruses (Haynes and Burton, 2017). While ˜50% of HIV-infected individuals generate bnAbs (Hraber et al., 2014), bnAbs in this setting only arise after years of infection (Bonsignori et al., 2016; Doria-Rose et al., 2014; Liao et al., 2013). BnAbs isolated from infected individuals have one or more unusual traits, including long third complementarity determining regions (CDR3s)(Yu and Guan, 2014), autoreactivity (Kelsoe and Haynes, 2017), large insertions and deletions (Kepler et al., 2014), and high frequencies of somatic mutations (Burton and Hangartner, 2016). Somatic hypermutation of the B cell receptor (BCR) heavy and light chain genes is the primary diversification method during antibody affinity maturation - the evolutionary process that drives antibody development after initial BCR rearrangement and leads to high affinity antigen recognition (Teng and Papavasiliou, 2007). Not all somatic mutations acquired during antibody maturation are necessary for bnAb development; rather high mutational levels may reflect the length of time required to elicit bnAbs (Georgiev et al., 2014; Jardine et al., 2016b). Therefore, shorter maturation pathways to neutralization breadth involving a critical subset of mutations is desirable because antibody mutation levels induced by vaccines seldom reach the mutation frequencies observed in bnAbs (Easterhoff et al., 2017; Moody et al., 2011). Importantly, within this subset of critical mutations some mutations may be probable and easy to elicit, while other mutations may be improbable and very challenging to induce due to biases in how mutations arise during affinity maturation.

Somatic hypermutation occurs prior to antigen affinity-based selection during affinity maturation (De Silva and Klein, 2015; Victora and Nussenzweig, 2012). The principal enzyme that mediates somatic hypermutation is activation-induced cytidine deaminase (AID) (Di Noia and Neuberger, 2007). AID preferentially targets specific nucleotide sequence motifs (referred to as “AID hot spots“) while targeting of other nucleotide motifs is disfavored (referred to as “AID cold spots“) (Betz et al., 1993; Pham et al., 2003; Yaari et al., 2013). AID initiates DNA lesions and their subsequent repair results in a bias for which bases are substituted at the targeted position (Cowell and Kepler, 2000). The consequence of this non-uniformly random mutation process is that specific amino acid substitutions occur with varying frequencies prior to antigenic selection. Mutations at AID hot spots can occur frequently in the absence of antigen selection due to immune activation-associated AID activity (Bonsignori et al., 2016; Yeap et al., 2015). Amino acid substitutions that occur infrequently generally require strong antigenic selection to arise during maturation (Brown et al., 1992; Kocks and Rajewsky, 1988). Such rare amino acid substitutions are improbable prior to selection for two reasons: 1) base mutations must occur at AID cold spots and 2) due to codon mapping, multiple base substitutions must occur for a specific amino acid change to take place. Within the critical subset of mutations that grant broad neutralization capacity to a bnAb lineage, those key mutations that are also improbable prior to selection may represent important events in bnAb maturation and are thus compelling targets for selection in a vaccine setting. We recently described a rare mutation in an early ancestor of a V3-glycan bnAb that allowed broad neutralization of transmitted/founder viruses demonstrating in one bnAb lineage that functionally important, improbable mutations can be roadblocks in HIV-1 bnAb development (Bonsignori et al., 2017). However, what has remained unclear is whether and to what extent such roadblocks are a general problem for bnAb elicitation. Here we describe the identification of improbable mutations in three bnAb B cell lineages, determine the functional relevance of these mutations for development of bnAb neutralization potency, and outline a vaccine design strategy for choosing sequential Envs capable of selecting B cells with BCRs with functionally important improbable mutations.

## RESULTS

### Identification of Functional Improbable Antibody Mutations

To determine the role of rare mutational events in bnAb development, we developed a computational program to identify improbable antibody mutations. Our program, <u>A</u>ntigen <u>R</u>eceptor <u>M</u>utation <u>A</u>nalyzer for <u>D</u>etection of <u>L</u>ow <u>L</u>ikelihood <u>O</u>ccurrences (ARMADiLLO) simulates the somatic hypermutation process using a model of AID targeting and base substitution via DNA repair (Yaari et al., 2013) and estimates the probability of any amino acid substitution in an antibody based on the frequencies observed in the computational simulation (**Figure 1**).

**Figure 1.**
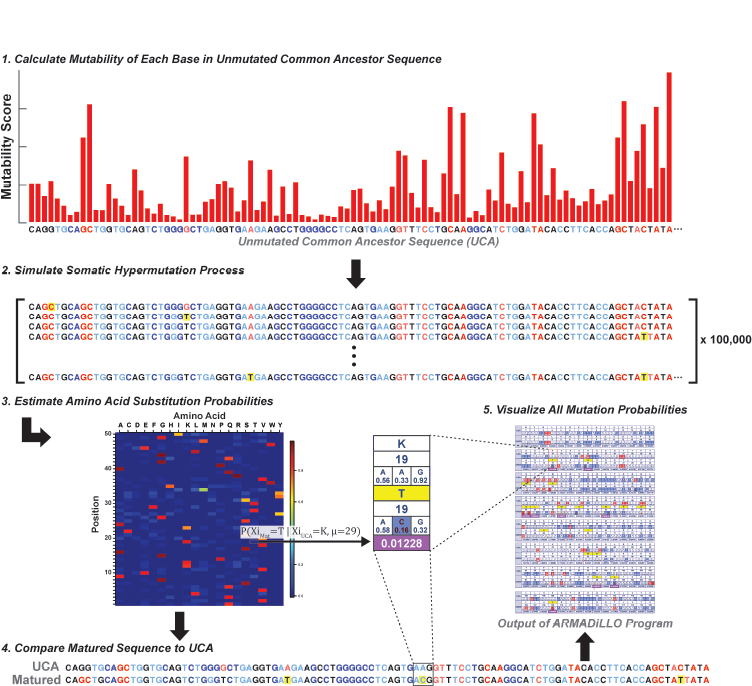
Computational method for estimating the probability of antibody mutations. The probability of an amino acid substitution during B cell maturation in the absence of selection is estimated by simulating the somatic hypermutation process. 1) The inferred unmutated common ancestor sequence (UCA) of the antibody of interest is assigned mutability scores according to a statistical model of AID targeting. 2) Bases in the sequence are then drawn randomly according to these scores and mutated according to a base substitution model (see methods). Rounds of single base mutation continue for the number of mutations observed in the antibody of interest with mutability scores updated as the simulation proceeds. The simulation is then repeated 100,000 times to generate a set of synthetic matured sequences. 3) An amino acid positional frequency matrix is constructed from the simulated sequences and utilized to estimate the probability of amino acid substitutions. 4) The UCA and matured sequence are aligned and 5) the estimated probability of amino acid substitutions identified in the matured sequence are outputted.

First, we applied ARMADiLLO retrospectively to the analysis of a mutation in a bnAb lineage that occurred at an AID cold spot that we have previously shown was functionally important for neutralization (Bonsignori et al., 2017). The DH270 V3-glycan bnAb lineage developed a variable heavy chain (V_H_) complementary determining region 2 (CDR H2) G57R mutation that when analyzed with the ARMADiLLO program was predicted to occur with <1% frequency prior to selection (**Figure S1**). This mutation was functionally critical because reversion back to G57 in the DH270 bnAb lineage resulted in total loss of neutralization potency and breadth (Bonsignori et al., 2017). Thus, the ARMADiLLO program can identify a known, key improbable mutation.

All BCR mutations arise during the stochastic process of somatic hypermutation prior to antigenic selection. In HIV-1 infection, antibody heterologous breadth is not directly selected for during bnAb development because BCRs only interact with autologous virus Envs. Since improbable bnAb mutations can confer heterologous breadth, they represent critical events in bnAb development, and make compelling targets for focusing selection with immunogens. To test this hypothesis, we analyzed three additional bnAb lineages with ARMADiLLO to identify improbable mutations (defined as <2% estimated probability of occurring prior to selection; see Materials and Methods) and then tested for the effect of these mutations on bnAb neutralization during bnAb B cell lineage development. We chose three lineages that allowed for study of different levels of maturation in bnAb development: CH235, mid-stage bnAb development (Bonsignori et al., 2017); VRC01, late stage bnAb development (Wu et al., 2015); and BF520.1, early stage bnAb development in an infant (Simonich et al., 2016).

### A Subset of Improbable Mutations Confer Heterologous Neutralization in CD4 Binding Site bnAb Lineages

CH235 is a CD4-binding site, CD4-mimicking (Gao et al., 2014) bnAb B cell lineage that evolved to 90% neutralization breadth and high potency over 5-6 years of infection and acquired 44 V_H_ amino acid mutations (Bonsignori et al., 2016). We identified improbable mutations in the heavy chain of an early intermediate member of the lineage (also termed CH235), reverted each to their respective germline-encoded amino acid, and then tested antibody mutants for neutralization against the heterologous, difficult-to-neutralize (tier 2) (Seaman et al., 2010) TRO.11 HIV-1 strain (**Figures 2A and S2A**). Single amino acid reversion mutations resulted in either a reduction or total loss of heterologous HIV-1 TRO.11 neutralization for each of three improbable mutations, K19T, W47L and G55W demonstrating that improbable mutations in the CH235 lineage were indeed critical and could confer heterologous neutralization.

**Figure 2.**
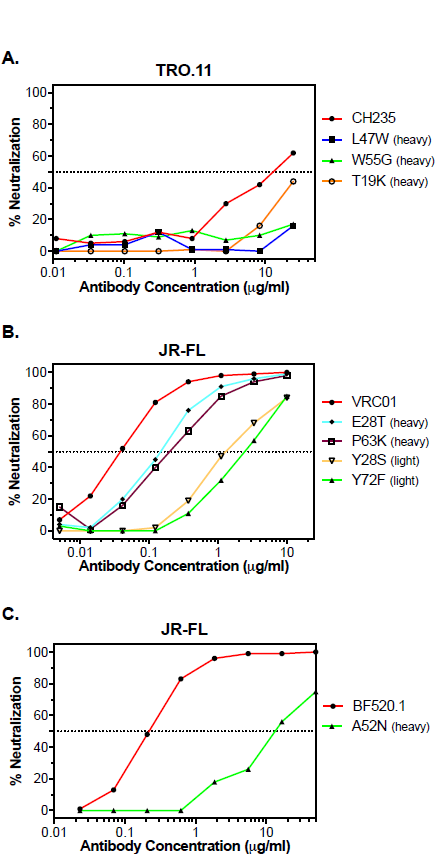
Improbable mutations confer heterologous neutralization in bnAb development. BnAbs A) CH235, B) VRC01 and C) BF520.1 and their corresponding mutants with reverted improbable mutations were tested for neutralization against heterologous viruses. The reversion of improbable mutations in all three bnAbs diminished neutralization potency.

Identification of the K19T mutation was of particular interest because the mutation was observed in all but one member of the CH235 bnAb lineage and was also present in two other CD4-binding site bnAbs (Scheid et al., 2011) from different individuals that shared the same VH gene segment (VH1-46) as CH235 (**Figure S3A**). We performed genomic sequencing of the individual from which the CH235 lineage was isolated and confirmed that K19T was a mutation and was not due to allelic variation in gene segment VH1-46 (**Figure S4**). Superposition of the CH235 complex into a fully-glycosylated trimer (Stewart-Jones et al., 2016) showed that the K19T mutation position was in close proximity to the N197 glycan site on the Env trimer (**Figures S3B and S3C**). The K19T mutation shortened the amino acid at this position thus allowing for larger glycan forms at the heterogeneously glycosylated N197 position (Behrens et al., 2016) providing a structural rationale for the effect of this mutation on heterologous breadth. Consistent with this hypothesis, CH235 neutralization of HIV-1 JR-FL, a tier 2 heterologous virus lacking the N197 glycan site, was unaffected by the T19K reversion mutation (**Table S1 and Figure S2A**). Moreover, we introduced the K19T mutation into the CH235 UCA and observed improved binding to an early autologous Env suggesting that the improbable K19T mutation may have been selected for by an early autologous virus variant (**Figure S3D**).

We next asked what role improbable mutations played in the maturation of a second broad and potent CD4 binding site-targeting bnAb lineage, termed VRC01, that acquired 43 V_H_ amino acid mutations (Zhou et al., 2010). We reverted improbable mutations in the VRC01 antibody and tested for their effects on neutralization of heterologous tier 2 HIV-1 JR-FL (**Figure 2B**). Reversion of improbable mutations reduced potency of heterologous neutralization of HIV-1 JRFL demonstrating that in the VRC01 CD4 binding site B cell lineage, single improbable amino acid substitutions can also have functional consequences for heterologous neutralization capacity. Improbable mutations identified by ARMADiLLO in the VRC01 light chain showed an even larger effect on reducing neutralization than heavy chain mutations (**Table S1 and Figure S2B**), further underscoring, along with an atypically short CDRL3 and a critical CDRL1 deletion (Zhou et al., 2013), the importance of key improbable events in the maturation of the VRC01 bnAb lineage.

### An Improbable Mutation Associated with Accelerated BnAb Development

Babies develop bnAbs earlier after HIV-1 infection than adults (Goo et al., 2014; Muenchhoff et al., 2016). We analyzed the glycan-V3 epitope targeting BF520.1 bnAb, isolated from an HIV-1 infected infant with many fewer mutations (12 V_H_ amino acid mutations) compared to VRC01 and CH235 bnAbs (Simonich et al., 2016). We identified an improbable mutation, N52A, located in the CDR H2 of BF520.1, reverted it to germline, and expressed the resultant antibody mutant (A52N). Heterologous neutralization of the A52N reversion mutant against tier 2 JR-FL virus was markedly reduced relative to wildtype BF520.1 (**Figure 2C**). The A52N reversion mutation antibody reduced neutralization potency for all tier 2 viruses that the BF520.1 bnAb could neutralize (**Table S1 and Figure S2C**) demonstrating that the N52A mutation was critical to the neutralization potency of BF520.1 and suggested the acquisition of this improbable mutation may have played a role in the relatively early elicitation (<15 months) of a bnAb with limited mutation frequency.

### Improbable Antibody Mutations Are Enriched in BnAbs

While not all improbable mutations will be critical for bnAb development, a subset will be key, as demonstrated in the examples above. To provide a view of the scope of the problem for many bnAb B cell lineages, we estimated the number of improbable mutations for a representative set of known bnAb lineages spanning all known sites of vulnerability on the Env trimer (**Figures 3A and S5, Table S2**). Study of a representative sample of bnAb lineages is plausible because of commonalities of Env recognition by bnAb germline precursors (Andrabi et al., 2015; Bonsignori et al., 2011; Gorman et al., 2016; Zhou et al., 2013). Compared to Env-reactive antibodies induced by an HIV-1 vaccine candidate (the RV144 vaccine) (Rerks-Ngarm et al., 2009) or antibodies isolated from non-HIV-1 infected individuals (Williams et al., 2015), the broadest and most potent HIV-1 bnAbs had the highest numbers of improbable mutations (**Figure 3B**). This result may follow directly from the observations that bnAbs tend to be highly mutated (**Figure S6A**) (Burton and Hangartner, 2016), and the number of improbable mutations an antibody possesses is correlated with its mutation frequency (**Figure S6B**) (Sheng et al., 2017). However, it is not known why most bnAbs are highly mutated. Recent work has shown that not all mutations in bnAbs are essential for neutralization activity (Jardine et al., 2016b). One hypothesis is that high mutation frequency is due to the extended number of rounds of somatic hypermutation required for a lineage to acquire a specific subset of mutations (Klein et al., 2013). If some of those specific mutations are also improbable, it is very likely that more probable mutations would be acquired prior to attaining key improbable ones. We found that for many bnAbs the number of improbable mutations exceeded what would be expected by chance given their high mutation frequency (**Table S3**). This observation, along with our experimental observations demonstrating that a subset of improbable mutations are important for neutralization capacity, is consistent with the notion that improbable mutations may act as key bottlenecks in the development of bnAb neutralization breadth. Thus, during chronic HIV-1 infection with persistent high viral loads that are required for bnAbs with improbable mutations to develop (Gray et al., 2011), excess numbers of probable mutations also accumulate. Probable mutations arise easily from the intrinsic mutability of antibody genes, and unlike improbable mutations, may not require Env selection (Bonsignori et al., 2016; Hwang et al., 2017; Neuberger et al., 1998). Thus, if the selection of critical improbable mutations can be targeted with Env immunogens, it should be possible to accelerate bnAb maturation and result in the induction of bnAb lineages with fewer mutations than those that occur in the setting of chronic HIV-1 infection.

**Figure 3.**
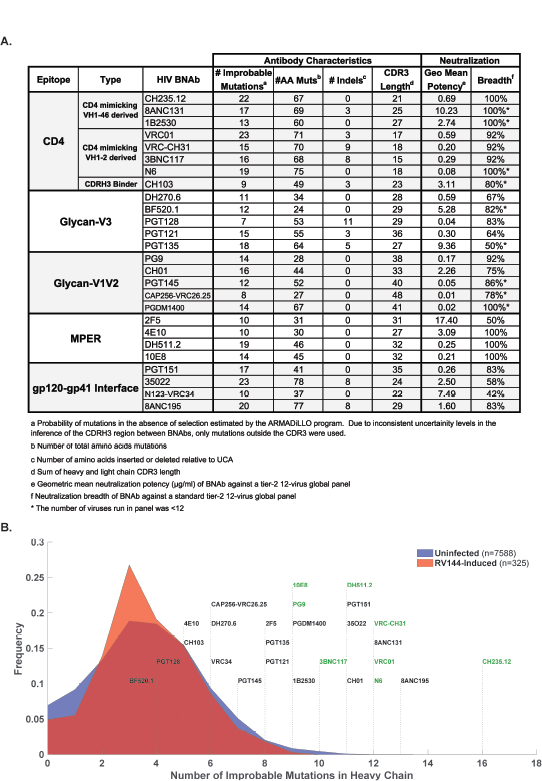
BnAbs are enriched for improbable antibody mutations. A) Table of improbable mutations for a representative set of bnAbs B) Histograms of the distributions of number of improbable mutations from antibody heavy chain sequences from three groups: “RV144-induced” antibodies were isolated from RV144 vaccinated subjects by antigenically sorting with RV144 immunogens (red shaded area); “Uninfected” antibodies correspond to duplicated NGS reads from IgG antibodies isolated from PBMC samples from 8 HIV-uninfected individuals (blue shaded area; see methods for details on sampling); a representative set of published bnAb antibody sequences are shown labeled above dotted lines that correspond to their number of improbable mutations (at the <2% level).

### Implications for Vaccine Design

The ability to identify functional improbable bnAb mutations using the ARMADiLLO program and antibody functional studies informs a mutation-guided vaccine design and immunization strategy (**Figure 4**). The principal goal is to be able to choose the correct sequential Envs to precisely focus selection towards the most difficult-to-induce mutations, while allowing the key easier, more probable mutations to occur due to antibody intrinsic mutability from immune activation-associated AID activity. In this strategy, improbable mutations are identified computationally using the ARMADiLLO program. Next, improbable mutations identified are expressed as single amino acid substitution mutant antibodies and their functional importance validated by Env binding and neutralization assays. Mutations in both maturation and reversion directions (UCA antibody with the single mutation added and mature antibody with the mutation reverted to germline, respectively) may be necessary to define the functional role of each mutation. Env mutants that bind to the UCA mutant and subsequent mutated intermediates with high affinity are chosen as immunogens to select for these functional improbable mutations. Last, sequential immunization with the chosen immunogens are studied for optimization of regimens to select for B cells with BCRs with the required improbable mutations. The vaccine strategy outlined here requires an adequate priming immunogen capable of engaging bnAb precursors (Dosenovic et al., 2015; Escolano et al., 2016; Jardine et al., 2016a; Jardine et al., 2015; McGuire et al., 2016; Sok et al., 2016; Williams et al., 2017; Zhang et al., 2016) prior to boosting to select for improbable mutations.

**Figure 4.**
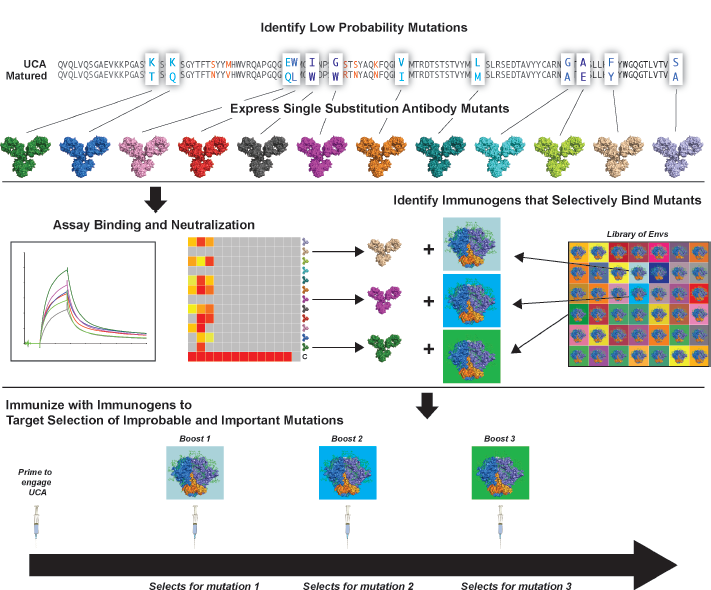
Mutation Guided Lineage Design Vaccine Strategy. Improbable mutations can act as important bottlenecks in the development of bnAbs and we propose here a strategy to specifically target those mutations for selection through vaccination. First, for a specific bnAb lineage, low probability mutations are identified computationally and recombinant antibody mutants corresponding to these mutations are produced (top panel). Binding and neutralization assays are performed to validate which of the improbable mutations are functionally important for lineage development (middle panel, left) and Envs are chosen that can specifically bind the corresponding antibody mutants (middle panel, right). These Envs are then used in a sequential immunization regimen to select the most difficult-to-induce, critical mutations thus potentially alleviating key bottlenecks in bnAb elicitation.

## DISCUSSION

Here we report the identification of improbable mutations in three bnAb lineages spanning multiple epitope specificities and representing distinct stages of bnAb development. We showed through functional antibody analysis that certain improbable mutations in these lineages were critical for the development of neutralization breadth and potency. In addition, we showed that bnAbs from HIV infected individuals were enriched for the number of improbable mutations relative to antibodies isolated from uninfected or HIV-1 vaccinated subjects. We conclude that improbable mutations that are functionally important for neutralization capacity can act as barriers for bnAb development in many bnAb lineages. Thus, the elicitation of important improbable mutations should be a high priority in vaccination. To this end, we proposed a vaccine design strategy that aims to target selection of critical improbable mutations with the goal of circumventing these barriers and accelerating the maturation of a bnAb response.

Vaccine strategies aiming to select for specific maturation mutations in a clonal lineage first requires adequate priming immunogens capable of engaging bnAb precursors. Recent progress has been made in the design of immunogens that bind bnAb UCAs (Jardine et al., 2016a; McGuire et al., 2013; Steichen et al., 2016; Zhang et al., 2016). Once UCAs are engaged, a vaccine must elicit mutations that keep antibody maturation on a pathway towards heterologous breadth. Immunization studies with fully germline-reverted bnAb UCA knock-in mice have so far failed to demonstrate the acquisition of the specific mutations that lead to full bnAb neutralization capacity (Briney et al., 2016; Tian et al., 2016) suggesting that the pathways that lead to breadth are blocked and not circumvented with the current generation of immunogens. We propose here that one significant barrier could be the acquisition of important mutations that are highly improbable and thus infrequent prior to selection in the germinal center.

If a mutation occurs prior to selection with less than 2% probability, the cutoff we use here to define an improbable mutation, its expected frequency in a germinal center would be less than 1 in 50 clonally related B cells. Estimates of the number of clonally related B cells range from 10-100 B cells per germinal center (Jacob et al., 1991; Tas et al., 2016) implying that the frequency of a single improbable mutation in any given germinal center will be two or fewer, per round of mutation. Thus, in order for improbable mutations to arise and be available for antigenic selection, bnAb B cell clones must either fortuitously acquire an important improbable mutation in the first round of mutation or must survive long enough in the germinal center to undergo many rounds of mutations and selection to increase the opportunities to acquire that improbable mutation. Moreover, B cell survival is dictated not just by having higher affinity to antigen than other clonally related members, but also by having higher affinity than other clones in the germinal center. In addition, heterologous neutralization breadth is never selected for directly and thus the competition is confounded by selecting for autologous antigen affinity. If an improbable mutation important for heterologous neutralization does not simultaneously increase affinity for the antigen present in the germinal center, it will not be selected. Thus, a vaccine design strategy must select for mutations that increase antibody affinity, however the sequence of acquisition of mutations is important and will require experimental determination. An additional caveat is that improbable mutations leading to different but chemically similar amino acids may be functionally equivalent.

Because improbable mutations arise as either rare intrinsic mutations or by selection by antigens derived from autologous virus, not all improbable mutations are required for mediation of heterologous neutralization (**Table S1**). Thus, the probability of a mutation is not predictive of its essential nature for neutralization breadth. While the recurrence of an improbable mutation within many clonal members can indicate positive selection, the selection criteria in B cell maturation is not heterologous neutralization capacity but rather is autologous antigen affinity. Consequently, improbable mutations must be evaluated for their functional relevance, and only the subset of improbable mutations that are also important for acquisition of neutralization breadth are considered potential bottlenecks for bnAb development. Thus, while we showed that bnAbs are enriched for the number of improbable mutations relative to antibodies from uninfected or vaccinated individuals, work remains to functionally assess all of the improbable mutations to define those required for bnAb affinity maturation. It is important to note as well that intrinsically mutable positions (Neuberger et al., 1998) can also be capable of conferring heterologous breadth. In this regard we identified one such functionally important probable intrinsic mutation in the CH235 lineage, S57R (**Table S1**). However, such highly probable mutations, by definition, are easily inducible and are not likely to represent barriers in bnAb development.

Interestingly, bnAbs that demonstrate relatively low numbers of improbable single somatic mutations (**Figure 3A**) possessed other unusual antibody characteristics that were due to additional improbable events such as insertion/deletions (indels) or extraordinary CDR H3 lengths. For example, the bnAbs with the two lowest number of improbable mutations were PGT128 and CAP256-VRC26.25. These bnAbs are notable for having the largest indels (PGT128; 11 aas) or the longest CDR H3 (CAP256-VRC26.25; 38 aas) of the bnAbs we analyzed. In summary, our data presented here suggest Env trimers evolved to evade neutralizing B-cell responses by hiding within low probability regions of antibody sequence space. The ARMADiLLO program and mutation-guided vaccine design strategy presented here should be broadly applicable for vaccine design for other mutating pathogens.

## Experimental Procedures

### Study Design

Malawian individual CH505, from whom the CH235 lineage was isolated and VH1-46 gene segment alleles determined by genomic sequencing, was enrolled in the Center for HIV/AIDS Vaccine Immunology (CHAVI) 001, nonblinded, nonrandomized, observational study protocol at a CHAVI clinical site in Malawi after informed consent was obtained under protocols approved by the Institutional Review Board of the Duke University Health System, the National Institutes of Health, and a clinical site review board in Malawi (Tomaras et al., 2011).

### Analysis of the Probability of Antibody Mutations

The probability of an amino acid substitution at any given position in the antibody sequence of an antibody of interest was estimated using the ARMADiLLO program. The algorithm and the analysis performed using ARMADiLLO are described in Supplemental Experimental Procedures.

### Antibody Site-directed Mutagenesis

BF520.1 mutant antibody genes were synthesized by Genscript and recombinantly produced. Mutations into antibody genes for CH235 and VRC01 mutants were introduced using the QuikChange II Lightning site-directed mutagenesis kit (Agilent Technologies) following the manufacturer‘s protocol. Single-colony sequencing was used to confirm the sequences of the mutant plasmid products. Primers used for introducing mutations are listed in the Supplemental Experimental Procedures.

### Recombinant Antibody Production

Antibodies were recombinantly produced as previously described (Saunders et al., 2017).

### HIV-1 Neutralization

Antibody neutralization was measured in TZM-bl cell-based neutralization assays as previously described (Li et al., 2005; Sarzotti-Kelsoe et al., 2014). CH235 and BF520.1 and selected mutants were assayed for neutralization using a global panel of 12 HIV-1 Env reference strains (deCamp et al., 2014). Neutralization values are reported as inhibitory concentrations of antibody in which 50% of virus was neutralized (IC_50_) with units in µg/ml.

### Antibody Binding Measurements

Binding of CH235.UCA and mutants to the monomeric CH505 transmitted/founder (T/F) delta7 gp120 and monomeric CH505 M5 (early autologous virus variant) delta8 gp120 (Bonsignori et al., 2016; Gao et al., 2014) was measured by surface plasmon resonance assays (SPR) on a Biacore S200 instrument and data analysis was performed with the S200 BIAevaluation software (Biacore/GE Healthcare) as previously described (Alam et al., 2013; Dennison et al., 2011).

## ACKNOWLEDGEMENTS

We would like to thank Bette Korber for many helpful discussions. We thank Shia-Mao Xia, Melissa Cooper, Kara Anasti, and Amanda Eaton for technical assistance. This work was funded by UM1 AI100645 from the Duke CHAVI– Immunogen Discovery, Division of AIDS, NIAID, NIH (to B.F.H.). T.B. and K.W. were supported by the Duke University Center for AIDS Research (CFAR), an NIH funded program (5P30 AI064518). R.R.M is supported by a Medical Scientist Training Program (MSTP) training grant (T32GM007171) and the Ruth L. Kirschstein National Research Service Award F30-AI122982-0, NIAID.

## SUPPLEMENTAL INFORMATION

**Figure S1.**
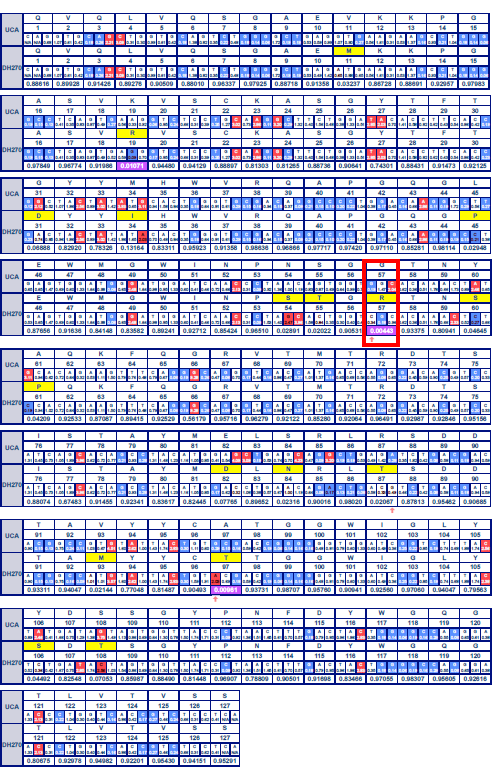
ARMADiLLO output for DH270 heavy chain shows G57R mutation is improbable. ARMADiLLO output for the DH270 heavy chain. The first three rows of each block corresponds to the sequence and the following four rows correspond to the matured DH270 sequenced. The first row is the amino acid sequence for the DH270 UCA. The second row is the amino acid numbering (consecutively numbered starting at 1 for the first residue) for the DH270 UCA. The third row is the sequence with each codon falling under the amino acid designated in row 1. The mutability score calculated with the S5F model is shown below the base in each box in this row. Each box is hot spots (red; mutability score>2) and cold spots (blue; mutability score <0.3). Row 8 is the estimated probability of the amino acid observed in the matured sequence (see methods for how this is calculated). The formatting pattern of rows 1-3 is repeated for the matured DH270 in rows 4-7. Amino acid substitutions are highlighted in yellow in row 4. Nucleotide mutations are shown in dark red text. mutations that are the result of mutations at AID cold spots are shown with an arrow below.

**Figure S2.**
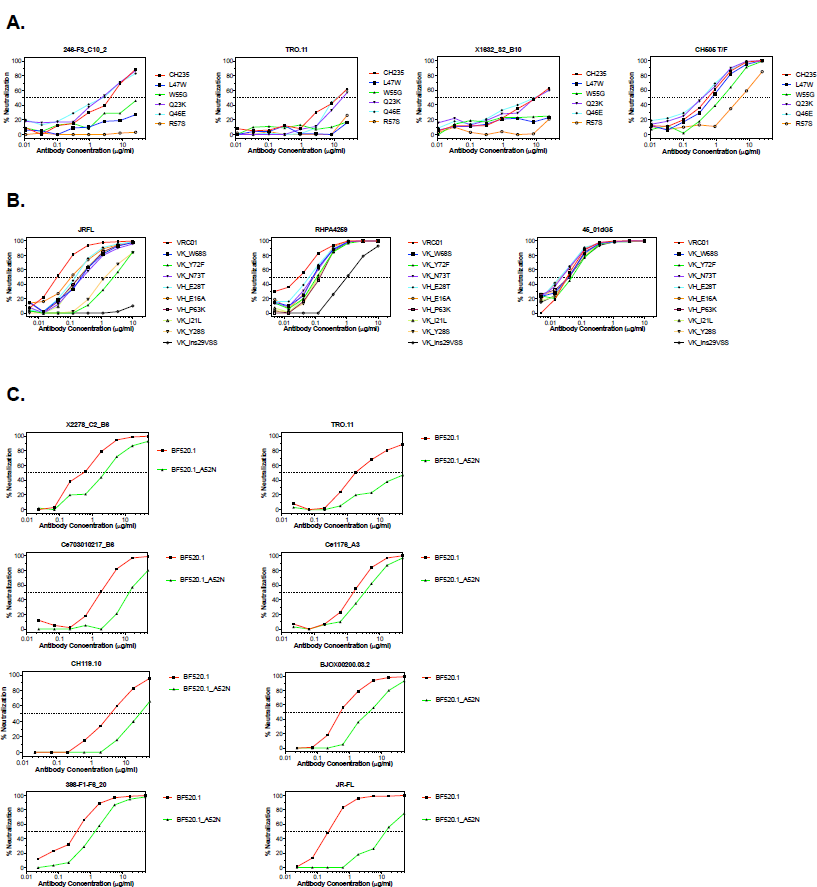
Neutralization of improbable mutation reversion mutants for CH235, VRC01, and BF520.1. Curves of the percent neutralization of WT (red line) **A)** CH235 **B)** VRC01 and **C)** BF520.1 and mutants containing reversions of identified improbable mutations against heterologous and autologous (CH505 T/F and 4501dG5 for CH235 and VRC01, respectively) viruses. 50% neutralization is denoted by a dotted line.

**Figure S3.**
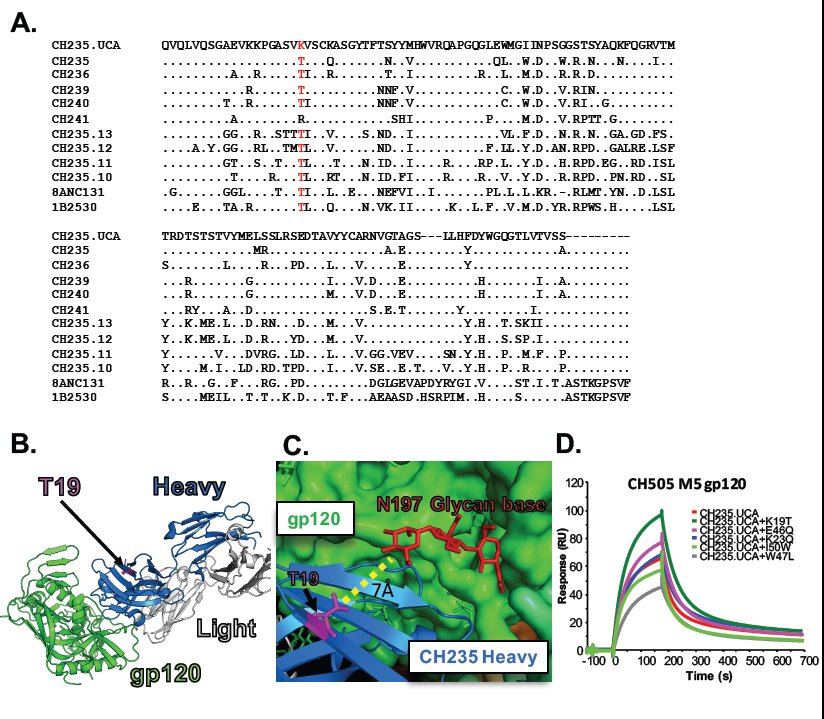
K19T mutation is conserved across all VH1-46 derived bnAb lineages and T19 position is glycan site. **A)** Amino acid multiple sequence alignment of the heavy chains of the three known VH1-46 gene segment-site bnAbs: 8ANC131, 1B2530, and the multiple member CH235 lineage aligned to UCA. The K19T mutation (red) is observed in all three lineages suggesting convergence of this three distinct individuals. Dots denote an amino acid match with the CH235 UCA in that **B)** The T19 position (magenta) in the CH235/gp120 complex structure (PDB: 5F9W) is outside of (heavy chain, blue; light chain, gray) binding site. The complex structure was determined with gp120 (green) and only minimal glycosylation (not shown) was resolved. **C)** Superposition of complex onto a fully glycosylated SOSIP trimer (5FYL) revealed that T19 (magenta) is in close (7Å) to the N197 glycan base (red) resolved in the trimer structure (green). A longer Lys residue position may sterically clash with longer glycans, providing a structural rationale for the the K19T mutation in VH1-46 derived CD4 binding site bnAbs. **D)** SPR sensorgrams for CH235 UCA and 5 UCA mutants containing improbable mutations show binding response to M5, construct featuring a single amino acid mutation from the CH505 T/F that makes it more favorable the CH235.UCA.

**Figure S4.**
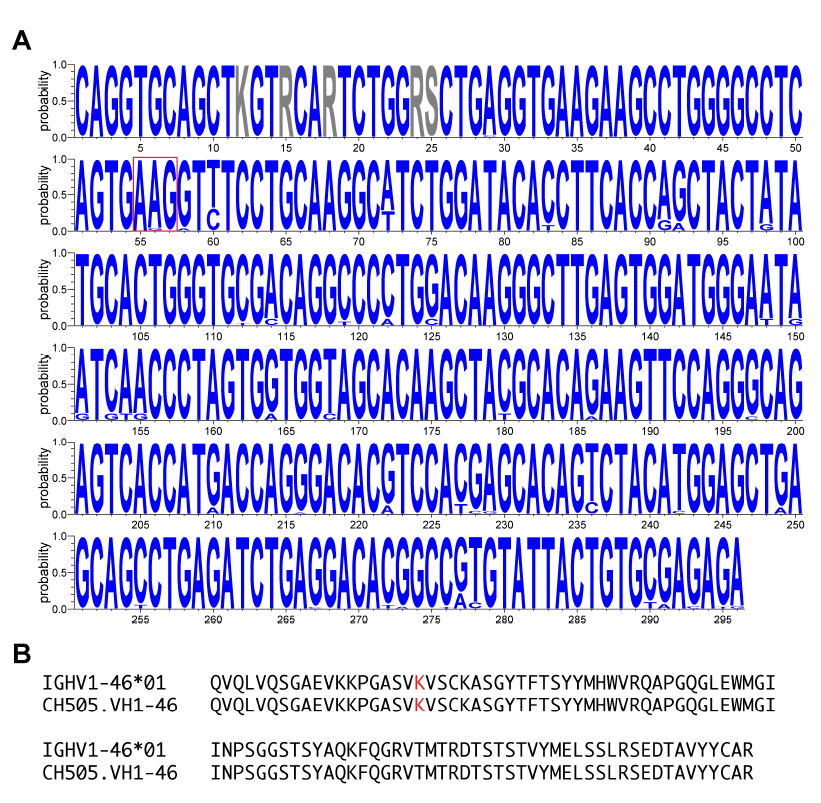
Genomic sequencing of the VH1-46 gene in the CH505 subject detected no allelic variation in the K19 codon confirming K19T was a mutation. **A)** Sequence logo plot of VH1-46 reads from genomic sequencing of the CH505 subject. No due to allelic variation were detected in the K19 codon (red box). Ambiguous bases in the are denoted in gray. **B)** Consensus of translated VH1-46 reads from the CH505 subject in amino acid sequence from the reference allele of IGHV1-46*01 confirming that a mutation and not due to an allelic variant.

**Table S1.** Neutralization of bnAbs and Mutants.

|  |  | Probability<sup>‡</sup> | CH505<br>T/F<sup>†</sup> | DU156.1<br>2 | JR-FL | Q23 | TRO.11* | 246-F3_C10<br>2* | X1632_<br>S2_B10* | 398-F1-F6_20* | CNE8* | X2278_<br>C2_B6* | CNE55* | CH119.1<br>0* | BJOX00<br>2000.03.<br>2* | 25710-2.43* | Ce70301<br>0217* | Ce1176_<br>A3* | SVA |
| --- | --- | --- | --- | --- | --- | --- | --- | --- | --- | --- | --- | --- | --- | --- | --- | --- | --- | --- | --- |
| CH235 | CH235 | - | 0.572 | 18.385 | 2.438 | 1.927 | 11.933 | 3.649 | 8.907 | >25 | >25 | >25 | >25 | >25 | >25 | >25 | >25 | >25 | >25 |
|  | T19K | 0.0116 | 0.888 | >50 | 1.635 | 7.39 | >50 | 6.148 | >25 | >25 | >25 | >25 | >25 | >25 | >25 | >25 | >25 | >25 | >25 |
|  | L47W | 0.00864 | 0.699 | NT | NT | NT | >25 | >25 | >25 | >25 | >25 | >25 | >25 | >25 | >25 | >25 | >25 | >25 | >25 |
|  | W55G | 0.00017 | 1.663 | NT | NT | NT | >25 | >25 | >25 | >25 | >25 | >25 | >25 | >25 | >25 | >25 | >25 | >25 | >25 |
|  | Q23K | 0.01612 | 0.398 | NT | NT | NT | 19.869 | 2.641 | 11.821 | >25 | >25 | >25 | >25 | >25 | >25 | >25 | >25 | >25 | >25 |
|  | Q46E | 0.01847 | 0.41 | NT | NT | NT | 12.142 | 2.178 | 7.125 | >25 | >25 | >25 | >25 | >25 | >25 | >25 | >25 | >25 | >25 |
|  | R57S | 0.22522 | 6.102 | NT | NT | NT | >25 | >25 | >25 | >25 | >25 | >25 | >25 | >25 | >25 | >25 | >25 | >25 | >25 |
|  | T19N | 0.033 | 23.249 | >50 | 20.289 | >50 | >50 | NT | NT | NT | NT | NT | NT | NT | NT | NT | NT | NT | >25 |
|  | T19R | 0.021 | 0.587 | 35.681 | 2.175 | 3.715 | 40.392 | NT | NT | NT | NT | NT | NT | NT | NT | NT | NT | NT | >25 |
|  | CH01+CH31 | - | <0.011 | 3.078 | <0.023 | <0.023 | 0.093 | 0.034 | <0.011 | 0.039 | 0.059 | 0.025 | 0.035 | 0.468 | 0.977 | 0.16 | <0.011 | 0.018 | >25 |

|  |  | Probability<sup>‡</sup> | JR-FL | TRO.11* | 246-F3_C10<br>2* | X1632_S<br>2_B10* | 398-F1-F6_20* | CNE8* | X2278_<br>C2_B6* | CNE55* | CH119.1<br>0* | BJOX00<br>2000.03.<br>2* | 25710-2.43* | Ce70301<br>0217* | Ce1176_<br>A3* | SVA |
| --- | --- | --- | --- | --- | --- | --- | --- | --- | --- | --- | --- | --- | --- | --- | --- | --- |
| BF520.1 | BF520.1 | - | 1.315 | 1.866 | NT | >50 | 0.39 | >50 | 0.467 | >50 | 3.894 | 0.598 | 40.449 | 1.901 | 1.472 | >50 |
|  | A52N | 0.0005 | 12.81 | >50 | NT | >50 | 1.247 | >50 | 2.238 | >50 | 22.203 | 3.971 | >50 | 13.288 | 3.634 | >50 |
|  | CH01+CH31 | - | 0.031 | 0.38 | NT | 0.078 | 0.108 | 0.183 | 0.055 | 0.113 | 2.29 | 10.806 | 0.895 | 0.102 | 0.11 | >50 |

|  |  | Probability<sup>‡</sup> | JR-FL | 45_01dG<br>5<sup>†</sup> | RHPA42<br>59 | SVA |
| --- | --- | --- | --- | --- | --- | --- |
| VRC01 | VRC01 | - | 0.04 | 0.036 | 0.033 | >10 |
|  | VK_W68S | 0.0007 | 0.215 | 0.038 | 0.089 | >10 |
|  | VK_Y72F | 0.009 | 2.095 | 0.053 | 0.109 | >10 |
|  | VK_N73T | 0.0085 | 0.261 | 0.025 | 0.088 | >10 |
|  | VH_E28T | 0.0009 | 0.141 | 0.019 | 0.069 | >10 |
|  | VH_E16A | 0.0012 | 0.12 | 0.022 | 0.092 | >10 |
|  | VH_P63K | 0.007 | 0.209 | 0.038 | 0.133 | >10 |
|  | VK_I21L | 0.0095 | 0.196 | 0.042 | 0.121 | >10 |
|  | VK_Y28S | 0.001 | 1.387 | 0.042 | 0.128 | >10 |
|  | VK_ins29VSS | ND | >10 | 0.042 | 1.054 | >10 |
|  | CH01+CH31 | - | <0.023 | <0.023 | 0.136 | >10 |

| Neutralization (IC50 µg/ml) |  |  |
| --- | --- | --- |
| 50 to 1 | 1 to .1 | <0.1 |

\*From global panel of 12 Tier-2 reference strains
† Autologous virus
‡ Estimated probability of mutation prior to selection as calculated by the ARMADiLLO program
NT Not Tested

**Figure S5.**
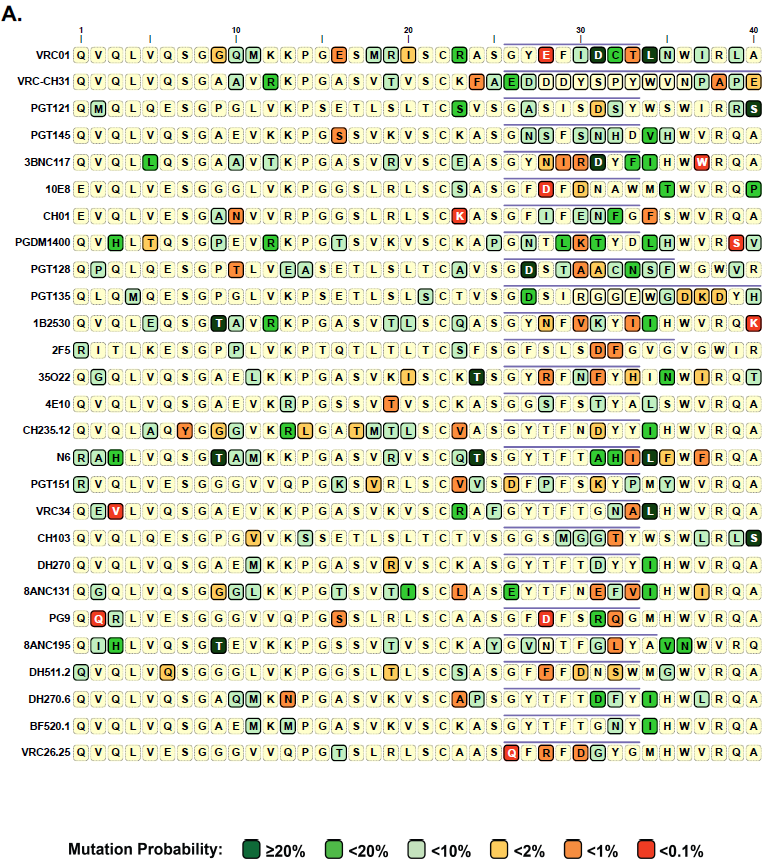
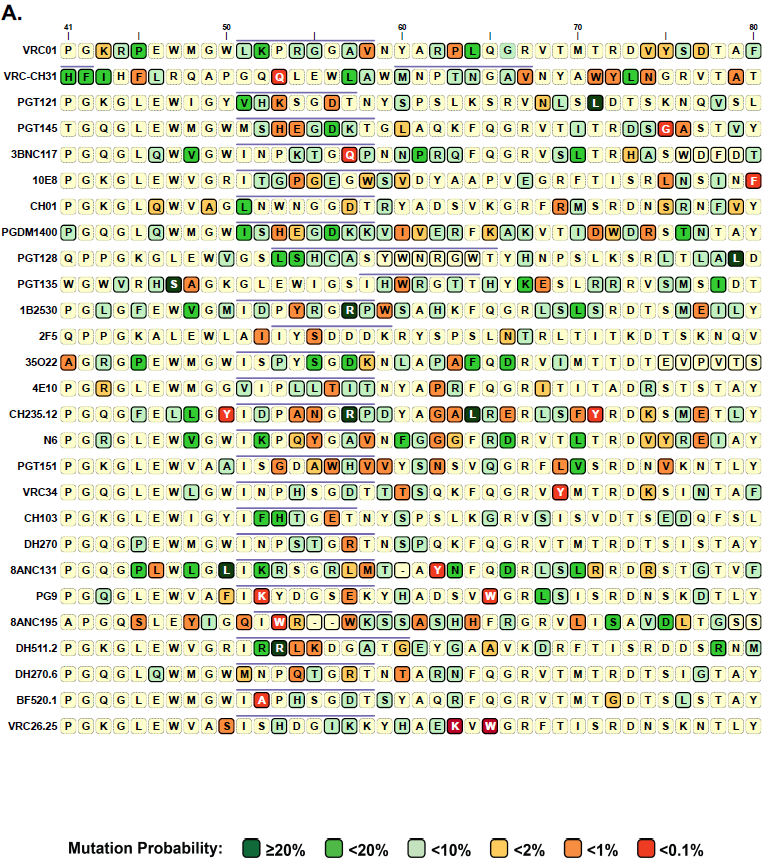
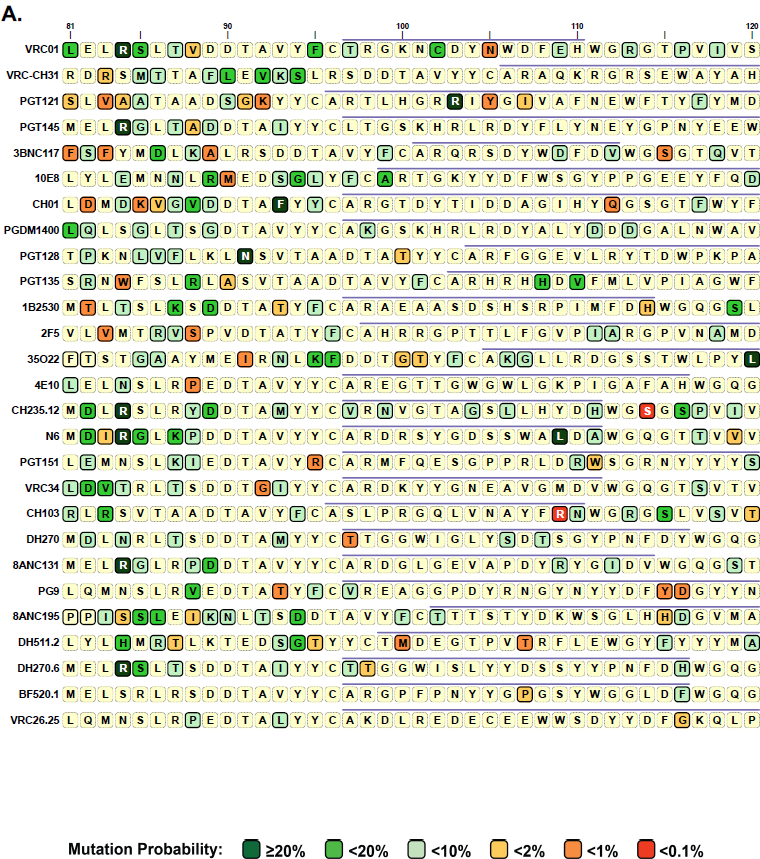
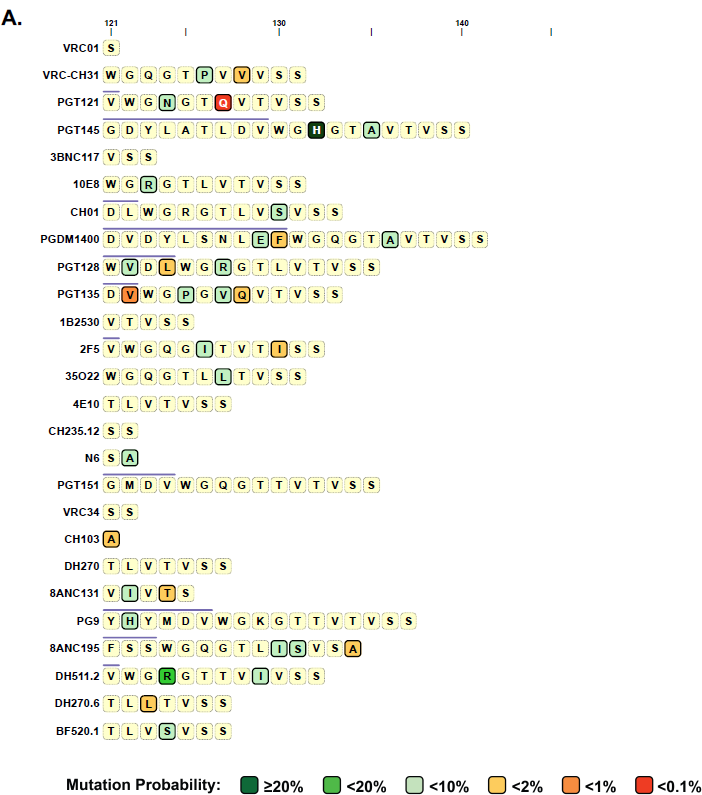
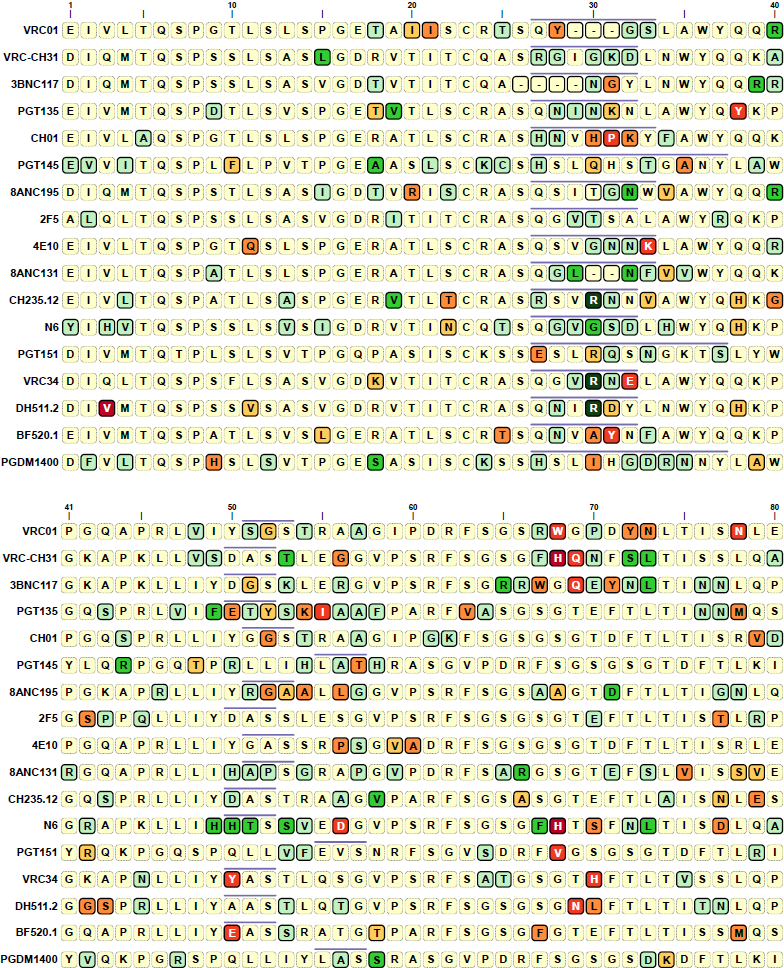
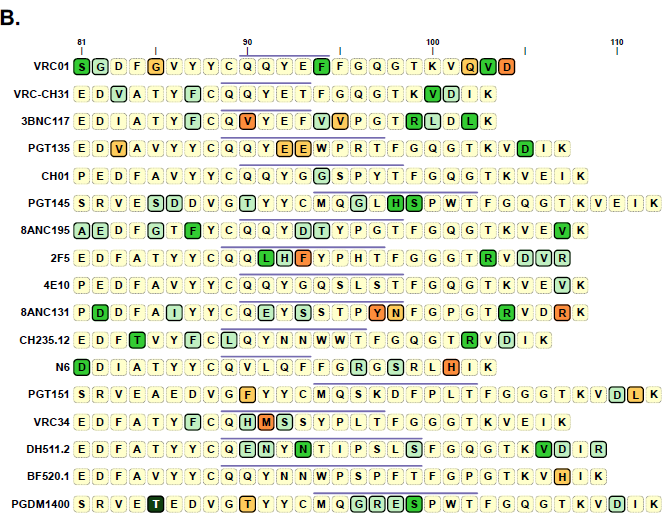
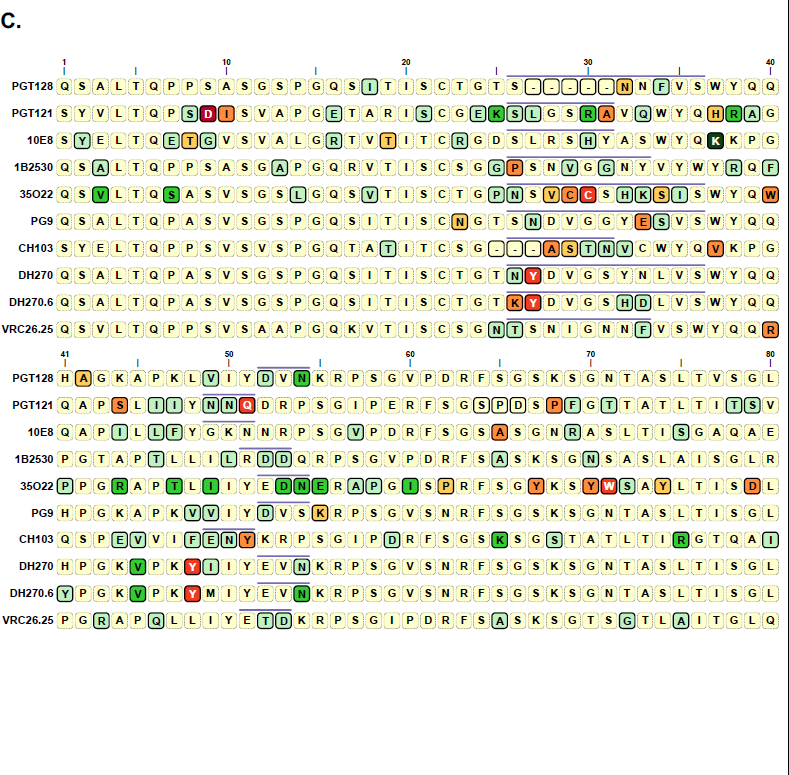
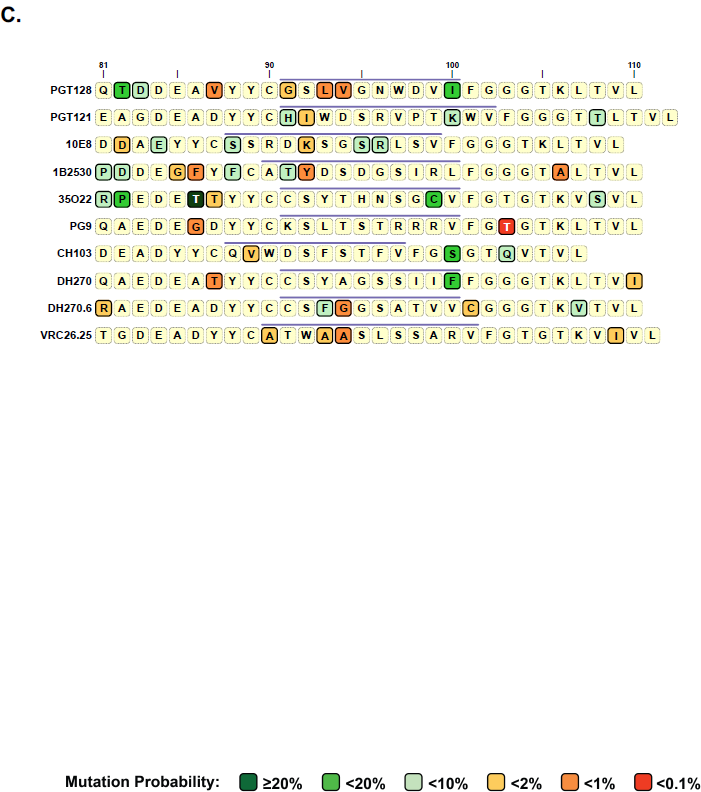
Representative bnAb sequences colored by mutation probability. A) <u>Heavy</u> chain sequences for a representative set of bnAbs are highlighted by their mutation probability ARMADiLLO. Mutations that are expected to occur frequently in the absence of selection (high probability mutations) are colored in shades of green (see legend). Mutations that are expected to rarely in the absence of selection (improbable mutations) are colored in shades of red (see legend). Amino acids residing in CDRs are denoted with a dark blue line above them. UCA inference was the observed bnAb sequence as input and as such there may be substantial uncertainty in mutation calls within the CDR3s. B) <u>Kappa</u> chain sequences for a representative set of bnAbs are highlighted by their mutation probability as estimated by ARMADiLLO. Mutations that are expected to occur frequently in the absence of selection (high probability mutations) are colored in shades of green (see legend). Mutations that are expected to rarely in the absence of selection (improbable mutations) are colored in shades of red (see legend). Amino acids residing in CDRs are denoted with a dark blue line above them. UCA inference was the observed bnAb sequence as input and as such there may be substantial uncertainty in mutation calls within the CDR3s. C) <u>Lambda</u> chain sequences for a representative set of bnAbs are highlighted by their mutation probability as estimated by ARMADiLLO. Mutations that are expected to occur frequently in the absence of selection (high probability mutations) are colored in shades of green (see legend). Mutations that are rarely in the absence of selection (improbable mutations) are colored in shades of red (see legend). Amino acids residing in CDRs are denoted with a dark blue line above them. UCA inference was performed with only the observed bnAb sequence as input and as such there may be substantial uncertainty in mutation calls within the CDR3s.

**Figure S6.**
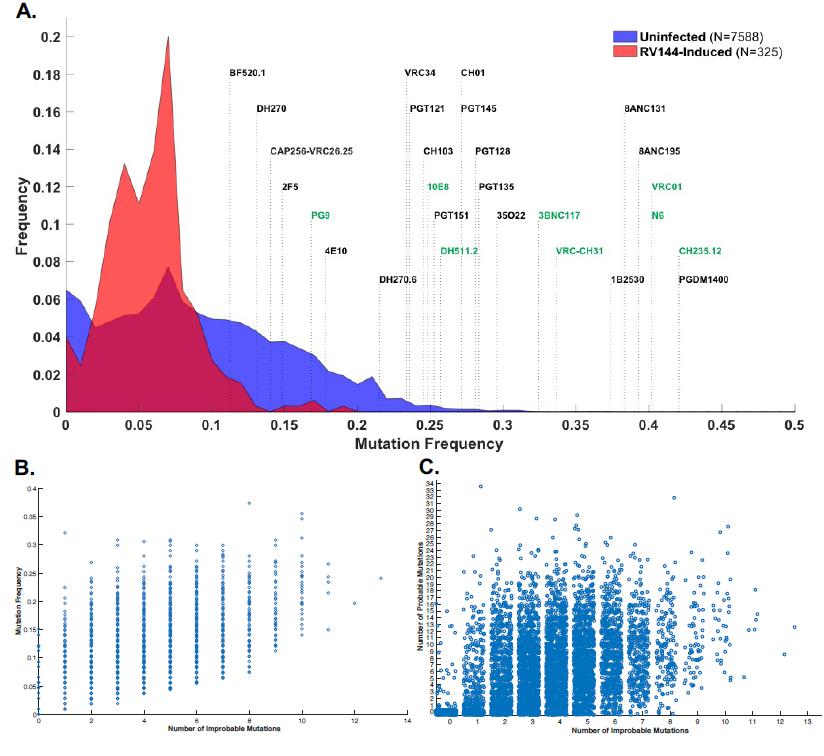
BnAbs have high mutation frequencies and mutation frequency is correlated with. **A)** Histograms for the distributions of number of improbable mutations (A) and mutation frequency (B) from heavy chain sequences from three groups: “RV144-induced” antibodies were isolated from RV144 vaccinated subjects by antigenically sorting with RV144 immunogens (red shaded area); “Uninfected” correspond to duplicated NGS reads from IgG antibodies isolated from PBMC samples from 8 individuals (blue shaded area; see methods for details on sampling); a representative set of bnAb antibody sequences are shown labeled above dotted lines that correspond to their mutation (defined as total number of amino acid mutations in non-CDRH3 VDJ sequence divided by non- sequence length). Scatterplots of **B)** number of improbable mutations versus amino acid frequency for 7588 NGS reads from uninfected IgG antibodies from PBMC samples from 8 HIV-individuals and **C)** number of improbable mutations versus number of probable mutations (≥2%). mutations was moderately correlated with number of probable mutations (Pearson‘s r=0.43). A stronger correlation was observed between improbable mutations and mutation frequency (Pearson‘s r=0.67) as expected because probable mutations are a subset of the total amino acid mutations amino acid mutation frequency. Jitter added in order to alleviate over-plotting in panel C.

**Table S3.** Number of improbable mutations expected by chance given mutation frequency.

| bnAb | #AA mutations<sup>A</sup> | AA mut freq | # Improbable Mutations (M)<sup>B</sup> | Expected Frequency of $\geq M^C$ | Counts |
| --- | --- | --- | --- | --- | --- |
| CH235.12* | 45 | 42.1% | 16 | 0.20% | 2/985 |
| 8ANC195* | 44 | 39.3% | 13 | 2.34% | 23/981 |
| N6* | 43 | 40.2% | 12 | 4.35% | 43/989 |
| VRC01* | 43 | 40.2% | 12 | 3.75% | 37/986 |
| 8ANC131* | 41 | 38.3% | 12 | 3.67% | 36/980 |
| VRC-CH31* | 39 | 33.6% | 12 | 1.43% | 14/981 |
| CH01* | 29 | 27.1% | 11 | 0.71% | 7/985 |
| DH511.2* | 28 | 25.7% | 11 | 0.51% | 5/981 |
| PGT151* | 27 | 25.2% | 11 | 0.62% | 6/974 |
| 35O22* | 34 | 29.6% | 11 | 1.32% | 13/984 |
| 3BNC117 | 36 | 32.4% | 10 | 5.28% | 52/985 |
| 10E8* | 27 | 24.8% | 9 | 4.54% | 45/991 |
| PGDM1400 | 45 | 42.1% | 9 | 36.59% | 360/984 |
| 1B2530 | 40 | 37.4% | 9 | 25.43% | 249/979 |
| PG9* | 18 | 16.8% | 9 | 1.63% | 16/983 |
| PGT135 | 32 | 28.3% | 8 | 15.96% | 157/984 |
| PGT121 | 25 | 23.6% | 8 | 8.61% | 85/987 |
| 2F5* | 16 | 14.8% | 8 | 4.47% | 44/984 |
| PGT145 | 29 | 27.1% | 7 | 25.96% | 256/986 |
| VRC34 | 25 | 23.4% | 6 | 32.21% | 315/978 |
| VRC26.25 | 15 | 14.0% | 6 | 23.71% | 234/987 |
| DH270.6 | 23 | 21.5% | 6 | 29.67% | 292/984 |
| CH103 | 26 | 24.5% | 5 | 56.58% | 559/988 |
| 4E10 | 19 | 17.8% | 5 | 46.97% | 465/990 |
| PGT128 | 32 | 28.1% | 4 | 82.36% | 808/981 |
| BF520.1 | 12 | 11.2% | 3 | 84.37% | 831/985 |

A Number of amino acid mutations, excluding CDRH3 region
B The number of improbable mutations M (at <2% cutoff) identified for the BNAb using ARMADILLO
C The expected frequency of M or greater improbable mutations (at <2% cutoff) from simulations of SHM at the BNAb's AA mutation frequency
\* Number of improbable mutations in this bnAb exceeds number expected by chance (cutoff=.05) given the mutation frequency of this bnAb

## SUPPLEMENTAL EXPERIMENTAL PROCEDURES

### Simulating the somatic hypermutation process

Because AID targets hot spots according to their underlying sequence motifs, the probability of mutations is sequence context dependent, making an analytical computation of the probability of a mutation in the absence of selection all but intractable. Instead, we take a numerical approach via simulation. Here, we estimate the probability of an amino acid substitution by simulating the somatic hypermutation (SHM) process and calculating the observed frequency of that substitution in the simulated sequences. The simulation proceeds as follows. Given a matured antibody nucleotide sequence, we first infer its unmutated common ancestor (UCA) sequence (Kepler, 2013; Kepler et al., 2014). Next, the matured antibody nucleotide sequence is aligned to the UCA nucleotide sequence and the number of sites mutated, t, is computed. Starting with the UCA sequence, 1) the mutability score of all consecutive sequence pentamers is computed according to the S5F mutability model (Yaari et al., 2013). 2) The mutability scores for each base position in the sequence are converted into the probability distribution, *Q*, by:

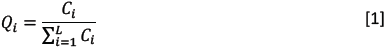

Where *C*_*i*_ is the mutability score at position *i* and *L* is the length of the sequence. 3) A base position, b, is drawn randomly according to *Q*. 4) The nucleotide n, at b, is substituted according to the S5F substitution model (Yaari et al., 2013), resulting in sequence *S*_*j*_ where j is the number of mutations accrued during the simulation. The procedure then iterates over steps 1-4 until j=t. This results in a simulated sequence, *S*_*t*_, that has acquired the same number of nucleotide mutations as observed in the matured antibody sequence of interest. If at any iteration during the simulation a mutation results in a stop codon, that sequence is discarded and the process restarts from the UCA sequence. This simulation procedure is then repeated to generate 100,000 simulated matured sequences. These nucleotide sequences are then translated to amino acid sequences.

### Estimating the probability of an amino acid substitution

The estimate of the probability of any amino acid substitution *U* → Y at site i given the number of mutations t observed in the matured sequence of interest is then calculated as the amino acid frequency observed at site i in the simulated sequences according to:

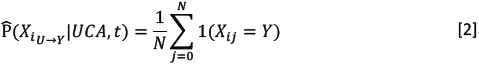

where *X*_*i*_ is the amino acid at site i which has the amino acid Y in the UCA sequence mutating to amino acid Y in the matured sequence of interest, UCA is the UCA sequence, N is the number of simulated sequences, 1 is an indicator function for observing amino acid Y at site i in the jth simulated sequence. This estimate is for an amino acid substitution *in the absence of selection* and we use this probability as a gauge of how likely it is that a B cell would arise to have this mutation prior to antigenic selection. Amino acid substitutions that are the result of mutations that occur in AID hot spots will have high probabilities, occur frequently and a subset of the reservoir of B cell clonal members would likely have these mutations present prior to antigenic selection. Amino substitutions that are the result of cold spot mutations or require multiple base substitutions will be much less frequent and could represent significant hurdles to lineage development and these substitutions may require strong antigenic selection to be acquired during B cell maturation.

### Improbable mutations

The probability of a specific amino substitution at any given position is the product of two components. The first component is due to the bias of the AID enzyme in targeting that specific base position and the DNA repair mechanisms preference for substituting to an alternative base. Practically speaking, substitutions that require mutations at AID cold spots and/or result in disfavored base substitutions by DNA repair mechanisms are infrequent and thus improbable. The second component is the number and length of available paths through codon space to go from an amino acid encoded by the codon in the UCA to that of the codon for the substituted amino acid in the matured sequence. To illustrate this, we turn to a practical example: the TAT codon which encodes the amino acid, Tyr. From the TAT codon, 5 amino acids are achievable by a single nucleotide base substitution (C,D,F,H,N,S), 12 amino acids by two base substitutions (A,E,G,I,K,L,P,Q,R,T,V,W) and 1 amino acid (M) by three base substitutions. Without considering the bias of AID, the Y->M mutation starting from the TAT codon is inherently unlikely to occur because it requires three independent mutational events to occur within the same codon. By simulating the SHM process, ARMADiLLO captures the interplay of these two components and is able to estimate the probability of any amino acid substitution prior to selection by taking both components into account.

For this study, we have selected a cutoff of less than 2% probability to classify mutations as “improbable“. We chose this cutoff to reflect a frequency in which the expected number of mutations in a B cell clone in a single germinal center would be less than or equal to 1. In order to calculate the expected number of mutations, we used estimates of the number of clonally related B cells in a germinal center. The range of these estimates is ˜10-100 clonally related B cells for immunizations in mice with various protein antigens (Jacob et al., 1991; Tas et al., 2016). Thus, for a mutation that has 2% probability, the expected number of B cells with this mutation in a germinal center is 1 in 50, reflective of one mutation per clone per germinal center. To demonstrate the effects of different choices of cutoffs, we applied additional cutoffs of 1%, 0.1% and 0.01% when calculating the estimated numbers of improbable mutations in a representative set of bnAbs and these additional data are included in an Excel spreadsheet in the supplementary materials.

### Calculating the expected number of improbable mutations

The number of improbable amino acid mutations, M, in an antibody sequence at a given probability cutoff can be estimated by applying [2] and enumerating over the entire amino acid sequence. For example, CH235.12 is estimated to have M=16 improbable mutations in its heavy chain when improbable mutations are defined as amino acid substitutions with <2% estimated probability. We estimate the probability of getting M improbable mutations or greater at a given amino acid mutation frequency, u, from the empirical distribution of the number of improbable mutations observed in sequences simulated to acquire T amino acid mutations, where T=u*L and L is the length of the sequence. To calculate the empirical distribution of improbable mutations for each antibody sequence of interest, we first randomly draw 1000 sequences from an antibody sequence dataset generated from NGS sequencing of 8 HIV-1 negative individuals and infer the UCA of each sequence (Kepler, 2013; Kepler et al., 2014). From these randomly sampled UCAs, we then simulate the SHM process using the same simulation procedure as detailed above and stop the simulation when each sequence acquires T amino acid mutations. This results in a set of 1000 simulated sequences each with an amino acid mutation frequency of u. The probability of observing M or greater improbable mutations in the absence of selection is then:

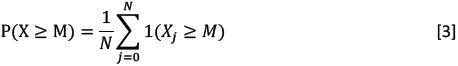

where N is the number of simulations (here N=1000), *X*_*j*_ is the number of improbable mutations in the jth simulated sequence (calculated from [2] over all amino acid positions in the sequence) and 1 is an indicator function. Here we exclude the CDR3 sequence from our calculations of both M and u as the inference of the UCA has widely varying levels of uncertainty in the CDR3 region depending on the input matured sequence.

Standard methods for determining selection at an amino acid site typically rely on the measure ω which is the ratio of non-synonymous mutations to synonymous mutations at that position in a multiple sequence alignment of related gene sequences. Here, we avoid this measure of selection for two reasons. In many instances in this study we have only two sequences to compare, the UCA and the matured sequence. This does not provide the number of observations needed for ω to reliably indicate selection. In some case, where we do have multiple clonal members to align, the number of mutational events at a site is also not sufficiently large enough for ω to be reliable. Secondly, ω is calculated under the assumption that non-synonymous mutations are of neutral fitness advantage. Clearly, due to the sequence dependence of AID targeting this assumption is violated in B cell evolution. Instead, we employ the heuristic that amino acid mutations that are estimated to be improbable yet occur frequently within a clone are likely to have been selected for. While indicative of selection, this too can be misleading if mutations occur early in a lineage, are neutral and generate a cold spot or *colder* spot, thus making it less likely for the position to mutate again. Thus, it is apparent that much work remains on developing rigorous methods for measuring selection in B cell evolution. Our approach here is to treat improbable amino acid mutations as candidates for selection and to ultimately confirm the fitness advantage conferred by such mutations through experimentally testing their effect on virus neutralization and antigen binding.

### Antibody sequences from HIV-1 negative subjects

We utilized a previously described next generation sequencing dataset generated from 8 HIV-1 negative individuals prior to vaccination (Williams et al., 2015). Briefly, to mitigate error introduced during the PCR amplification, we split the RNA sample into two samples, A and B, and performed PCR amplification on each, independently. Only VDJ sequences that duplicated identically in A and B were then retained. This approach allowed us to be highly confident that nucleotide variations from germline gene segments that occurred in the NGS reads were mutations and not error introduced during PCR. We refer to this dataset as “uninfected“.

### Antibody sequences from RV144-vaccinated subjects

We utilized a previously described set of antibody sequences (Easterhoff et al., 2017) isolated from subjects enrolled in the RV144 HIV-1 vaccination trial (Rerks-Ngarm et al., 2009). Antibody sequences were isolated from peripheral blood mononuclear cells (PBMC) from 7 RV144-vaccinated subjects that were antigen-specific single-cell sorted with fluorophore-labeled AE.A244 gp120 d11 (Liao et al., 2013). We refer to this dataset as “RV144-immunized“.

### Analysis of Improbable Mutations in BnAbs

Sequences of HIV-1 bnAbs were obtained either from NCBI GenBank or from the bNAber database (Eroshkin et al., 2014). For the comparison of improbable mutations for the representative set of bnAbs, improbable mutations were calculated using the ARMADiLLO program described above. UCAs were inferred using Cloanalyst (Kepler, 2013; Kepler et al., 2014). While many bnAbs had multiple clonal lineage member sequences available, some bnAbs had no other members isolated. Because of this, only the single sequence of the matured bnAb was used in the UCA inference in order to provide equal treatment of all sequences. Because uncertainty in the UCA inference is highest for the bases in the CDR3 region, precise determination of some mutations in this region is not feasible and we therefore ignored the CDR3 region in our analysis of the representative set of bnAbs. In the simulations, we prohibited any mutations from occurring in the CDR3 region by setting the probability of AID targeting to 0 for each base in the CDR3. Neutralization data for the bnAbs was obtained through the CATNAP database (Yoon et al., 2015) and corresponds to neutralization in the global panel of 12 HIV-1 Env reference strains (deCamp et al., 2014). For the calculation of geometric mean neutralization, undetectable neutralization was set to 100 µg/ml. Breadth was reported for all viruses that were tested and for several bnAbs (8ANC131, 1B2530, N6, CH103, BF520.1, PGT135, PGT145, VRC26.25, PGDM1400) neutralization data was not available for all 12 viruses in the global panel.

### Genomic sequencing of the VH1-46 gene segment

To confirm that K19T was a mutation and not an allelic variant in subject CH505 from which CH235 clonal members were isolated, we sequenced CH505 IGHV gene segments according to a previously described experimental protocol for high throughput genomic sequencing of Ig gene segments which is detailed in Scheepers et al (Scheepers et al., 2015). Sequencing was performed using the Illumina MiSeq platform. A custom sequence analysis pipeline was used to analyze the sequencing data for identifying novel alleles. Forward and reverse reads were merged using FLASh (Magoc and Salzberg, 2011) and quality filtered using the FASTx toolkit (http://hannonlab.cshl.edu/fastx_toolkit/). Primers were trimmed such that only the V region of the merged read was retained and all resulting reads were then de-duplicated. All sequences with fewer than 10 reads were discarded. We then aligned all sequences to all known IGHV gene segments using a custom semi-global pairwise alignment program. Sequences that matched closest to any reference VH1-46 allele (01-03) were retained. Sequences that included stop codons, reflective of potential pseudogenes, were discarded. We then built a multiple sequence alignment of the remaining sequences and produced a sequence logo plot weighted by the number of read copies for each sequence. For the 19^th^ codon, 96% of reads matched the VH1-46*01 reference AAG triplet encoding a lysine, consistent with homozygous lysine at this position (the remaining 4% of reads are consistent with the error level introduced during PCR amplification). The sequencing thus confirms K19T in the CH235 clone was indeed a mutation in the CH505 subject. A consensus of the translated reads was identical to the entire VH1-46*01 reference allele demonstrating no non-synonymous polymorphisms indicating that at the protein level the CH505 subject was homozygous VH1-46*01.

### Antibody Site-directed Mutagenesis Primers

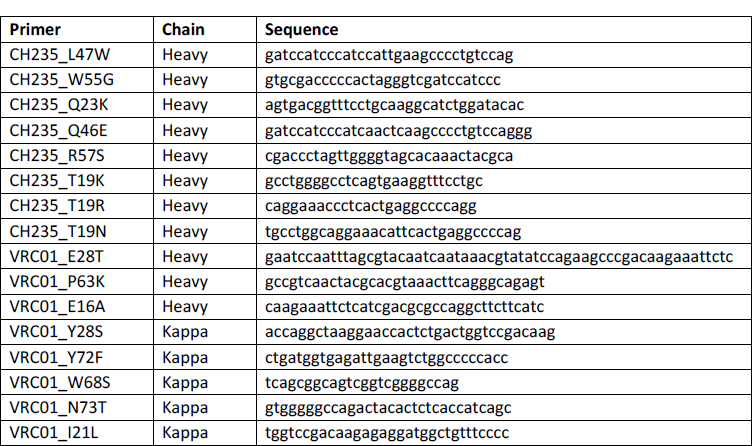

